# Electroadhesion of polymer networks by polycation interfacial bridging: sticky electrophoresis, ionic complexation, and chain entanglement

**DOI:** 10.64898/2026.06.05.730541

**Authors:** Binbin Ying, Kun-Hao Yu, Shu Yang, Jiawei Yang

## Abstract

An e-GLUE is a polymer network containing interpenetrating polycations, which can bond the anionic network of mucosa through interfacial polycation bridging under an electric field. Such an electroadhesion involves electrophoresis of polycations, ionic complexation between polycations and the anionic network, and polycation-network entanglement, yet their quantitative understanding is lacking. Here, we formulate a theoretical model to describe electroadhesion of polymer networks by polycation interfacial bridging. We use a diffusion-drift model coupled with a Bell-like field-dependent chain friction to describe the sticky electrophoresis of polycations in an anionic sea. The formation of ionic bonds is determined by local availability of cations and anions over the penetration depth. To debond, a force must either pull polycations out from the e-GLUE network or first dissociate them from ionic complexes and then pull out from the anionic network. We model chain pullout from the bulk networks to the interface as a viscous drag against water. The adhesion strength is calculated by summing the debonding force for each polycation per unit area across all chains. Our model quantitatively links electric field strength, applied duration, polycation chain length, and cation concentration to polycation electrophoresis kinetics, ionic bond formation, and adhesion strength. We further conduct electroadhesion tests, and our model predicts well with the experimental data. Lastly, we discuss the use of the model to guide the e-GLUE design.

**TOC graphic:** 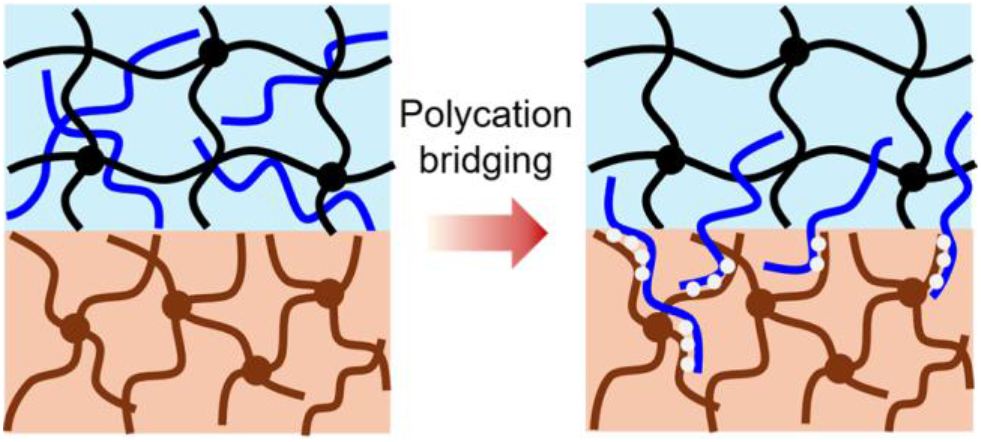

For Table of Contents use only

## 1. Introduction

Mucosa can be seen as an anionic polymer network, as one of its major components is anionic proteins^1^. Hydrogel adhesion to mucosa with prolonged retention is challenging because continuous motility, food passage, and rapid tissue turnover deteriorate adhesion^2, 3^. Traditional mucoadhesives^4^ and hydrogel adhesives^5, 6^ form physical or chemical bonds to the superficial mucosa, which can be cleared out in less than 24 hours. We have proposed that adhesion to mucosa over a certain depth rather than on the surface can prolong retention and have thereby developed an e-GLUE—a neutral polymer network containing interpenetrating polycations—to achieve and retain strong mucosal adhesion up to 30 days^7^. When the e-GLUE and the mucosa are brought into contact and subject to an electrical field, the polycations migrate across the interface to the mucosa by active electrophoresis and passive diffusion (Fig. 1a) and then ionically complex with mucosa anionic proteins (Fig. 1b). As a result, those polycations lying across the interface, with a part of the chain segments in physical entanglement with the polymer network of the e-GLUE, and the other part in ionic complex and physical entanglement with the mucosa anionic network, establish adhesion. The polycations that either have not reached the mucosa or have been completely in the mucosa cannot bridge the interface to contribute to adhesion.

**Fig. 1.**
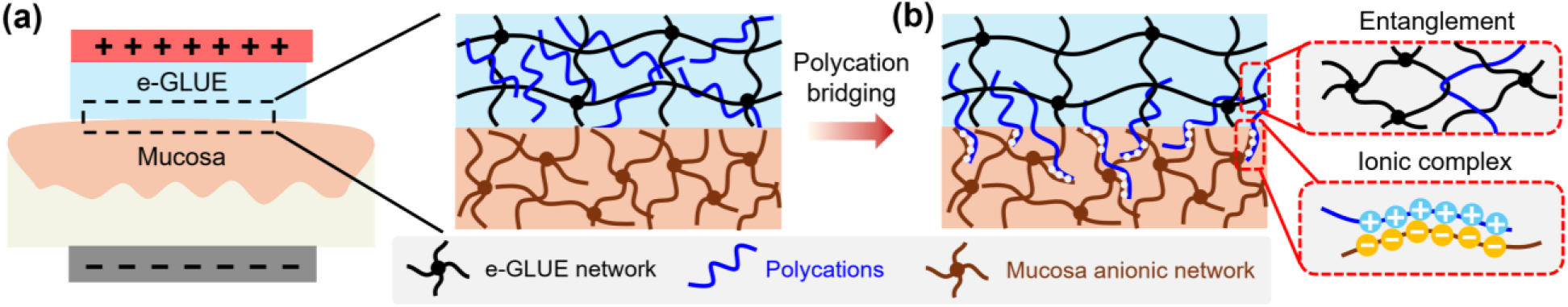
Electroadhesion by polycation interface bridging. **(a)** Illustration of an e-GLUE of a polymer network containing polycations, and a mucosa of an anionic polymer network. **(b)** Under an electric field, the polycations penetrate the mucosa and subsequently form ionic complexes. Polycations with a part of the chain segments in physical entanglement with the polymer network of the e-GLUE, and the other part in ionic complex and physical entanglement with the mucosa anionic network, bridge the interface.

An electric field has been applied to bond hydrogels and tissues. For example, a cationic hydrogel and an anionic hydrogel (e.g., tissue) can ionically complex at the interface under an electric field, and they can be dissociated under a reversed field^8, 9^. Alternatively, a hydrogel containing interpenetrating polycations and another hydrogel containing interpenetrating polyanions can bond through electrophoresis of polycations and polyanions toward the interface and formation of complexes and entanglements^10, 11^. Unlike these systems, ours is established between a neutral hydrogel containing interpenetrating polycations and an anionic hydrogel.

To measure adhesion strength, probe-pull tests are often performed, in which a perpendicular tensile force is applied to detach two adhered materials, and the adhesion strength is calculated by the maximum force divided by the adhesion area^12-14^. When the e-GLUE and the anionic network are detached, the interface-bridging polycations must be pulled out from either network. Pulling out from the e-GLUE network side requires overcoming the friction along the pulling path, whereas pulling out from the anionic network needs to first rupture the ionic complexes. The adhesion strength is the force to pull out the bridging chains, either from the e-GLUE or the anionic network, per unit area. Nevertheless, a quantitative model describing the supramolecular process and the formation of adhesion is lacking, which is important to provide a design principle for the e-GLUE to guide, tailor, and optimize adhesion strength and kinetics.

In this paper, we present a theoretical model together with experiments to quantitatively study electroadhesion by polycation interfacial bridging. We describe chain electrophoresis by a diffusion-drift model, coupled with a Bell-like field-dependent chain friction, which gives the polycation distribution in the anionic network with time. The polycations must penetrate smaller than a critical depth, beyond which they no longer bridge the interface and do not contribute to adhesion. The number of ionic bonds that can be formed is determined by the local availability of cations and anions within the penetration depth. Upon debonding, polycations are pulled out from either network. We model this by viscous drag against water. We finally determine the adhesion strength by summing the smaller force to pull out a chain per unit area over all chains. We further conduct electroadhesion tests to validate the model. This work provides a theoretical foundation for a range of polymer transport and interaction behaviors under an electric field.

The paper is organized as follows. Section 2 sets up the model. Section 3 develops the model and determines the adhesion strength. Section 4 presents model results. Section 5 describes electroadhesion experiments and results, and compares them with theoretical predictions. Section 6 discusses the use of these findings for e-GLUE design. Section 7 gives the conclusion.

## 2. Model setup

In the e-GLUE, polycations are physically entangled within the neutral polymer network of the hydrogel. Each polycationic chain can be seen as confined in a tube due to surrounding entangled chains (Fig. 2a). Under an electric field, electrostatic forces are generated on the cationic units of the chain, which drive the chain to migrate toward the anode (Fig. 2b) The chain migrates in a reptation motion under combined diffusion and electrophoretic drift along a curvilinear path in the tube (Fig. 2c). When the cationic unit of the chain encounters the anionic site, an ionic pair is formed. We point out that due to counterion competition, steric exclusion, and geometric mismatch, ion complexation can occur at any site of the chain, and both chain ends can remain mobile^15, 16^. Given that a maximum electric field applied in experiments is 8,000 V/m and a cationic unit carries one electric charge of 1.6 × 10^−19^ C, the electrostatic force on the cationic unit is ∼10^−15^ N, which by itself cannot break the ionic pair (force is on the order of 10^−11^ N)^17^. Nonetheless, the ionic pair is dynamic and reversible, which can dissociate with the current pair and reassociate with a neighboring ionic site to form a new pair^18-20^. Such a pair exchange is local and often fast compared to the long-range chain migration. Therefore, ionic bonds do not immobilize a chain but act as stickers that retard chain migration^21, 22^. In addition, solvent drag and topological constraints can also resist migration. The diffusivity for polymer migration in a polymer network typically ranges from 10^−11^ to 10^−13^ m^2^/s, depending on the chain length^23-25^, while the diffusivity for sticky migration can be as low as 10^−20^ m^2^/s (ref^18, 20^).

**Fig. 2.**
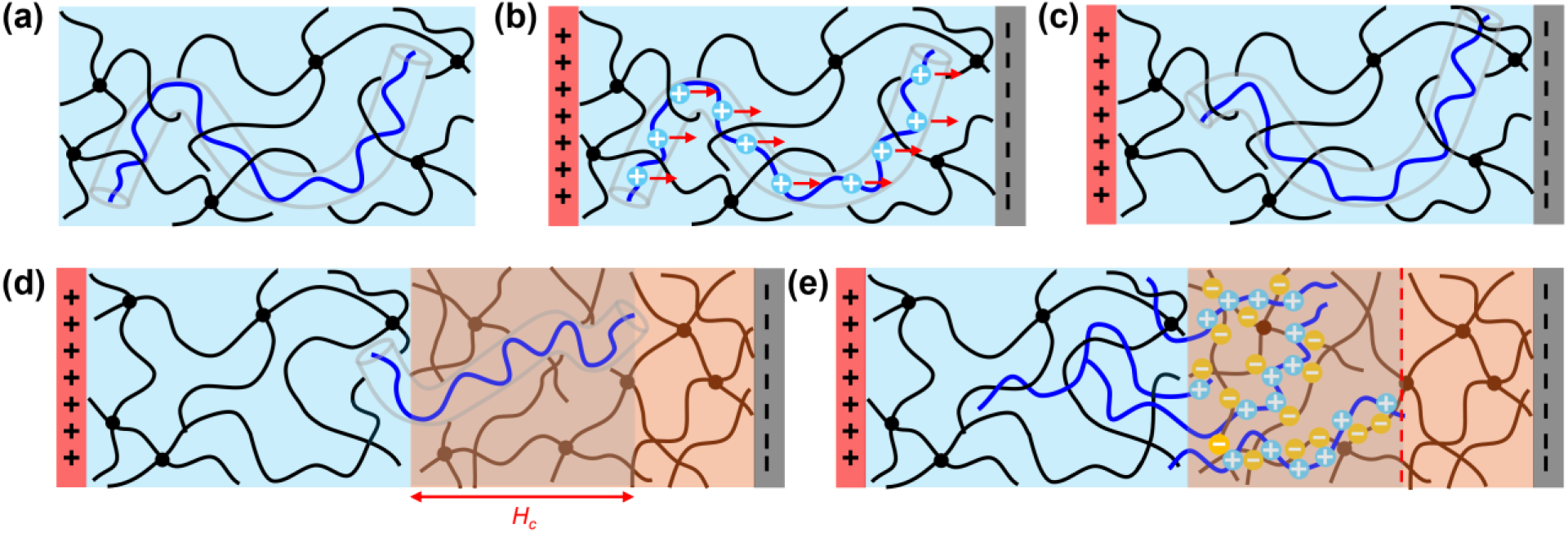
Electrophoresis, ionic complexation, and entanglement in electroadhesion. **(a)** A polycation is initially confined in a tube. **(b)** Under an electric field, the electrostatic forces (red arrows) drive the chain to migrate towards the anionic network. **(c)** The chain migrates via reptation. **(d)** The chain penetrates the anionic network and migrates through it in a sticky reptation motion. **(e)** The chains bridging the interface enable adhesion.

To bridge the interface, the chain must occupy a certain depth in both networks. A critical penetration depth *H*_*c*_ is on the order of the end-to-end distance of the chain, beyond which the chain is completely in the anionic network and can no longer bridge the interface (Fig. 2d). A longer chain gives a bigger critical penetration depth. The subsequent chains continuously migrate in and progressively complex with remaining anionic sites (Fig. 2e). Note that not every cation can pair with an anion to form an ionic bond, and its location and the number along the chain are uncertain and most likely different from each other. To simplify analysis, we introduce a bond formation parameter, assuming ionic bonds are formed uniformly along the entire chain, yielding a statistical average of ionic bonds per chain.

Upon debonding, the interface-bridging polycations are pulled out. Since the force to break a covalent bond is much higher than that to break ionic complexes, pulling a chain out from the e-GLUE network would require overcoming the viscous drag along the chain; the force scales with the contour length of the chain. Pulling a chain out from the anionic network first breaks ionic bonds and then overcomes the viscous drag along the chain. The force scales with the number of ionic bonds and the contour length of the chain in the anionic network.

Which network a chain is pulled out from depends on the relative force requirement between the two competing networks. Consider three states of chain-bridging with a penetration depth *H*: (i) a chain that just penetrates the anionic network, *H* → 0, (ii) a chain that penetrates all the way to the critical depth, *H* → *H*_*c*_, and (iii) a chain with 0 < *H* < *H*_*c*_ (Fig. 3a). In state (i), the chain is extensively entangled with the e-GLUE network, which is more likely to be pulled out from the anionic network. In state (ii), the chain is both ionically bonded and entangled with the anionic network, which is more likely to be pulled out from the e-GLUE network. In state (iii), a penetration depth *H*_*p*_ of equal chain pullout exists, at which the force that pulls out polycations from the e-GLUE network is the same as that ruptures the ionic bonds and pulls out polycations from the anionic network. When 0 < *H* < *H*_*p*_ (e.g., state (i)), the chains are pulled out from the anionic network; when *H*_*p*_ < *H* < *H*_*c*_ (e.g., state (ii)), the chains are pulled out from the e-GLUE (Fig. 3b). Therefore, the adhesion strength contributed from the chain pullout can be determined by summing the pullout force of a chain per unit area over all chains across the interface.

**Fig. 3.**
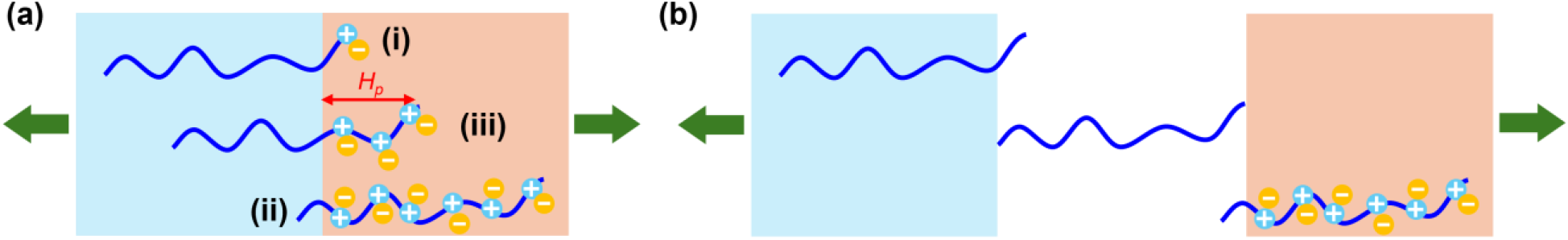
Debonding mechanism. **(a)** Three representative states of chain-bridging. A penetration depth *H*_*p*_ exists at which the force that pulls the chain out from either network is the same. **(b)** Chain pullout from which network depends on penetration depth.

## 3. A theoretical model

### 3.1. Electrophoresis of polycations

For simplicity, we model polycations as monodisperse, consisting of *n* Kuhn segments of length *b*. The contour length of the chain is *l*_*c*_ = *bn*, and the end-to-end distance of the chain in the tube is *l*_*e*_ = *bn*^1/2^. An electric field *E* pushes the chain in the global *y* direction, but the chain migrates along the curvilinear *s* coordinate (Fig. 4a). The projection of the electric force onto the *s* coordinate contributes to chain migration. Here, *s* represents the statistical average of migration paths across all chains, as each chain follows its own random walk^26-28^. As shown later, the *s* coordinate can be mapped to the global *y* coordinate (Fig. 4b).

**Fig. 4.**
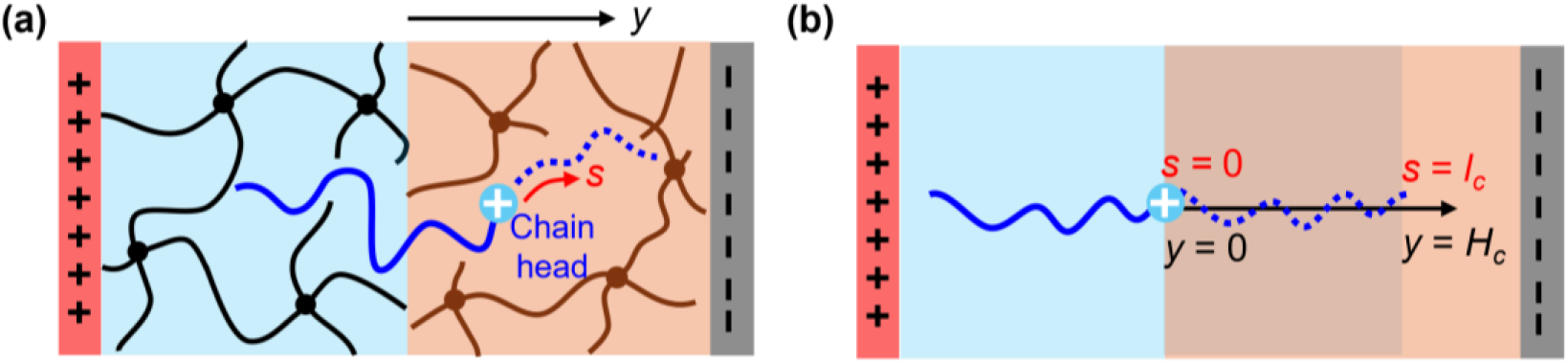
Local and global coordinates in chain migration. **(a)** An electric field aligns with the global *y* coordinate, whereas a chain migrates along the curvilinear *s* coordinate. **(b)** 1D map between the *s* coordinate and the *y* coordinate.

Because a chain spans multiple locations, we track the position of the chain head, a distal Kuhn segment at the chain’s front, to identify the entire chain’s location. Or equivalently, a chain is treated as a particle, and once the particle location is found, the segments of an entire chain trail the particle. Let the molar concentration of chain heads at a coordinate *s* and at a time *t* be *C*(*s, t*) (mol/m^3^) and the molar flux of chain heads be *J* (mol/m^2^·s). Conservation of mass gives

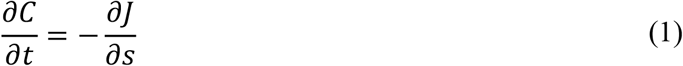

The flux consists of concurrent polymer diffusion and electrophoretic drift. Adopting the Stokes-Einstein description of molecular friction^16, 29-31^,

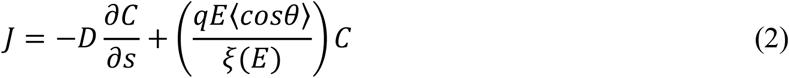

where *D* = *k*_*B*_*T*/*nξ*(*E*) is the Rouse diffusivity^32^, *q* is the charge per Kuhn segment, *k*_*B*_ is the Boltzmann constant, *T* is the absolute temperature, *E* is the electric field, and *ξ*(*E*) is the electric-field-dependent friction coefficient per Kuhn segment and is negatively correlated with the magnitude of the electric field. A longer chain and a larger friction lead to a slower migration. The factor <cos*θ*> represents the average projection of the electric field onto the local tangent of the *s* coordinate. For isotropic tube orientation, this orientation factor can be approximated as^16, 30, 31^,

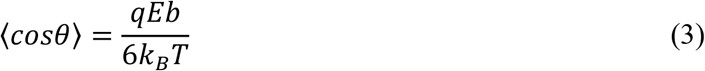

Eq. (3) can be understood as follows. The drift velocity of a Kuhn segment along the *y* direction is *v*_*drift*_ = *μE*, where *μ* is the electrophoresis mobility. The diffusion velocity of a Kuhn segment along the *s* coordinate is *v*_*diff*_ ∼ *D*/*b*. The orientation factor can be estimated as *v*_*drift*_/*v*_*diff*_ ∼ *μEb/D*. Using the Nernst-Einstein equation to relate diffusivity to electrophoresis mobility and carried charges, *D* = *k*_*B*_*Tμ*/*q*, gives *v*_*drift*_/*v*_*diff*_ ∼ *qEb/k*_*B*_*T*. Combining Eqs. (1) – (3) yields the coupled diffusion-drift equation for chain migration,

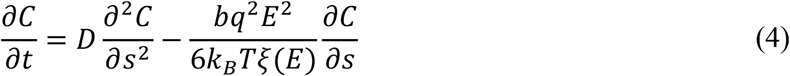

The first term on the right is chain diffusion, and the second term is chain electrophoresis. *C*(*s, t*) describes the spatiotemporal distribution of chain heads in the anionic network.

Because the chains within the critical penetration depth, or the chain contour length, can contribute to adhesion, we solve Eq. (4) in the domain 0 ≤ *s* ≤ *l*_*c*_. At *s* = 0, the chain head just crosses the interface; At *s* = *l*_*c*_, the chain head reaches *H*_*c*_ (Fig. 4b). At *t* = 0, no polycation is in the domain, and the initial condition is

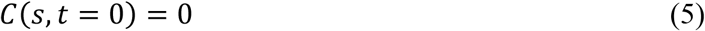

Assuming the e-GLUE is a reservoir of polycations in a short time, which maintains a constant concentration of chain heads at *s* = 0. The boundary condition is

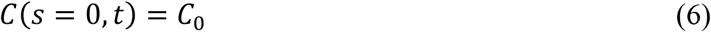

where *C*_0_ is the molar concentration of polycations in the e-GLUE in the preparation. The corresponding molar concentration of cations *C*_0_^*cat*^ = *mnC*_0_, where *m* is the number of monomers in one Kuhn segment, and *mn* is the degree of polymerization. At *s* = *l*_*c*_, we impose an open-outflow condition that allows chains to exit the domain. Eq. (4) can be solved numerically with Eq. (5) and (6).

### 3.2. Electric-field-assisted reduction of chain friction

To capture that chain migration accelerates with increasing electric field^7^, we introduce a field-dependent friction coefficient per Kuhn segment by a Bell-like activation model^33-35^,

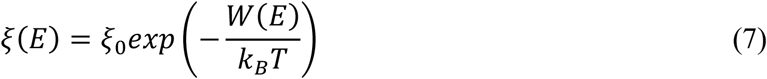

where *ξ*_0_ is the zero-field friction coefficient and *W*(*E*) is an activation energy, which can be expressed as the work done by the electric field to displace a chain over an activation length *L*,

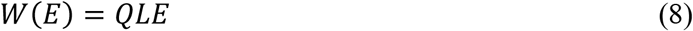

Because the ionic pair exchange is local, the relevant local length is the Kuhn length, i.e., *L* ∼ *b*. The exact value needs to be determined by fitting experimental data. *Q* is the total charge of the chain, as we treat a chain as a particle in migration, i.e., *Q* = *qn*. This model has been applied to cell adhesion^36^, mussel adhesion^37^, single-molecule tests^38, 39^, cationic-anionic bond formation^16^, and protein folding^40^.

### 3.3. Formation of ionic complexes

During polycation migration, when the local cation concentration is lower than the anion concentration, the ionic bond concentration is limited by the cation concentration. Conversely, when the local cation concentration becomes higher than the anion concentration, e.g., given a sufficiently long time, the ionic bond concentration is limited by the anion concentration. Because the anion concentration is a constant, when polycations occupy all anions, the ionic bond concentration plateaus.

To quantitatively capture this trend, we need to calculate the cation concentration at each location during migration. At a location *s* and at time *t*, the cations include the chain heads that migrate exactly to *s*, and those chain segments at *s*, but their chain heads have already passed *s* but not reached *l*_*c*_. This is equivalent to finding all chain heads in the domain between *s* and *l*_*c*_ and placing them at *s* in a domain of one Kuhn length. Because one Kuhn segment carries *m* cations, the molar concentration of cations at *s* can be written as

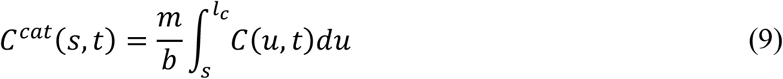

We map the *s* coordinate to the global *y* coordinate by the treatment adopted in polymer interdiffusion and healing models^26, 27, 35^,

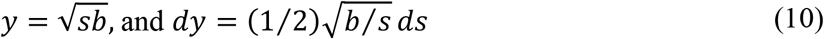

When *s* = *l*_*c*_ = *bn*, the critical penetration depth is

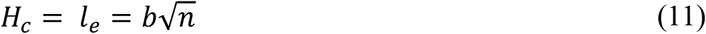

Substituting Eq. (10) into Eq. (9), *C*^*cat*^(*s, t*) can be converted to *C*^*cat*^(*y, t*). In the following, we compare the local concentrations of cations and anions in the *y* coordinate.

The anionic network carries one anion in every monomer and is uniformly distributed in space, independent of time. The molar concentration of anions is

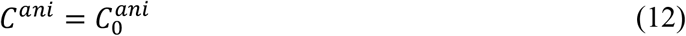

At a penetration depth *y*, the concentration of ionic bonds is determined by the local availability of cations and anions, namely,

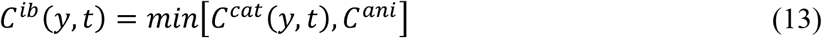

Here, the function *min*[*α, β*] means choosing the smaller value between *α* and *β*. Equivalently, this condition can be written piecewise as

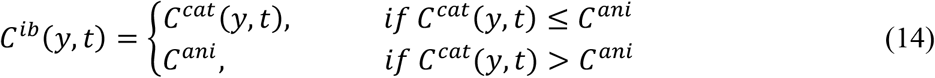

### 3.4. Stress for breaking ionic complexes

As discussed in Section 2, debonding breaks ionic bonds when the chain penetration depth is smaller than *H*_*p*_, corresponding to the contour length *s*_*p*_ in the *s* coordinate (Fig. 3, state (i)). Using Eq. (10), *H*_*p*_, the projection of *s*_*p*_ to the *y* direction, can be written as

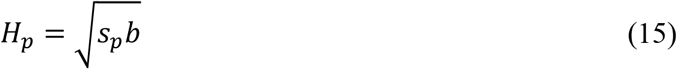

We note that Eq. (9) describes the total cation distribution in the anionic network, which includes cations on the chains in both domains within *H*_*p*_ and beyond *H*_*p*_. Yet, only those cations that belong to chains in the domain within *H*_*p*_ contribute to ionic bonds that break during debonding. The molar concentration of these cations is written as

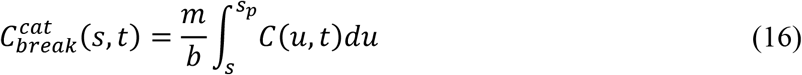

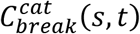 can also be converted to 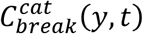 by Eq. (10). To calculate the ionic bonds that break during debonding, we need to rewrite Eq. (13) to only account for the contribution of 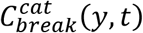. Because all cations have an equal opportunity to form ionic bonds, we assume proportional occupation of anions according to the ratio of 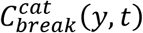 and 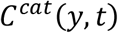. Combining Eq. (9), (13), and (16) gives the molar concentration of ionic bonds that break during debonding,

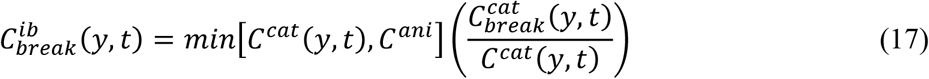

Eq. (17) gives the ideal case of ionic bond formation in which every cation can pair with an anion. We introduce a bond formation parameter *λ* between 0 and 1 to account for the real number of ionic bonds formed along the chain. Here, *λ* = 0 means no bond is formed, and *λ* = 1 means a complete formation of bonds. Using Eq. (17) to account for chains within the domain *H*_*p*_, the stress to break ionic bonds is

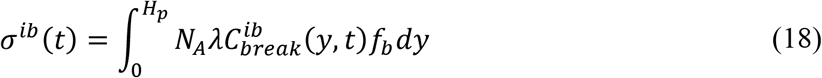

where *N*_*A*_ is Avogadro’s number and *f*_*b*_ is the force to break an ionic bond.

### 3.5. Stress for chain pullout

A force pulling a single chain out against water can be estimated by the Stokes drag,

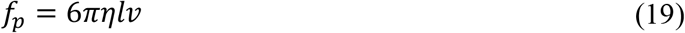

where *l* is the chain contour length being pulled out, *η* is the viscosity of water, and *v* is the pullout velocity. When a chain has its contour length *s* penetrated in the anionic network, *l* = *s*, the contour length in the e-GLUE network is *l* = *l*_*c*_ – *s*. The equal chain pullout contour length *s*_*p*_ can be obtained by setting the force to pull the chain out of the e-GLUE network and the anionic network equal,

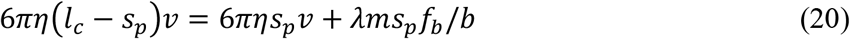

which gives

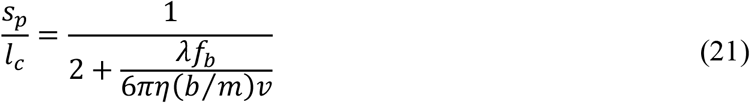

Here, *b*/*m* is the monomer length. The competition between the force to break an ionic bond *λf*_*b*_ and the force to pull the chain over a monomer distance 6π*η*(*b*/*m*)*v* determines *s*_*p*_. When *λ* = 0, no ionic bond forms, and *s*_*p*_ = 0.5*l*_*c*_, which ensures pulling the chain out from either network is equal. As *λ* becomes bigger until 1, more ionic bonds form, and the chain penetrates less to make the equal pullout, *s*_*p*_ < 0.5*l*_*c*_ (Fig. 5a). Plug in typical values (listed in Fig. 5) and set *λ* = 0.1, *λf*_*b*_/6π*η*(*b*/*m*)*v* ∼ 100 and *s*_*p*_ = 0.011*l*_*c*_, implying that just a fraction of the chain entering the anionic network is already sufficient to reach the equal chain pullout condition. Under fast pulling, *v* → ∞, an infinitely large drag force is generated, and the chain cannot be disentangled, equivalently as if they are covalently bonded. In this case, the force to break an ionic bond becomes negligible, and *s*_*p*_ = 0.5*l*_*c*_. Under slow pulling, *v* → 0, the drag force is zero, and any formation of ionic bonds cannot balance the force, *s*_*p*_ = 0 (Fig. 5b).

**Fig. 5.**
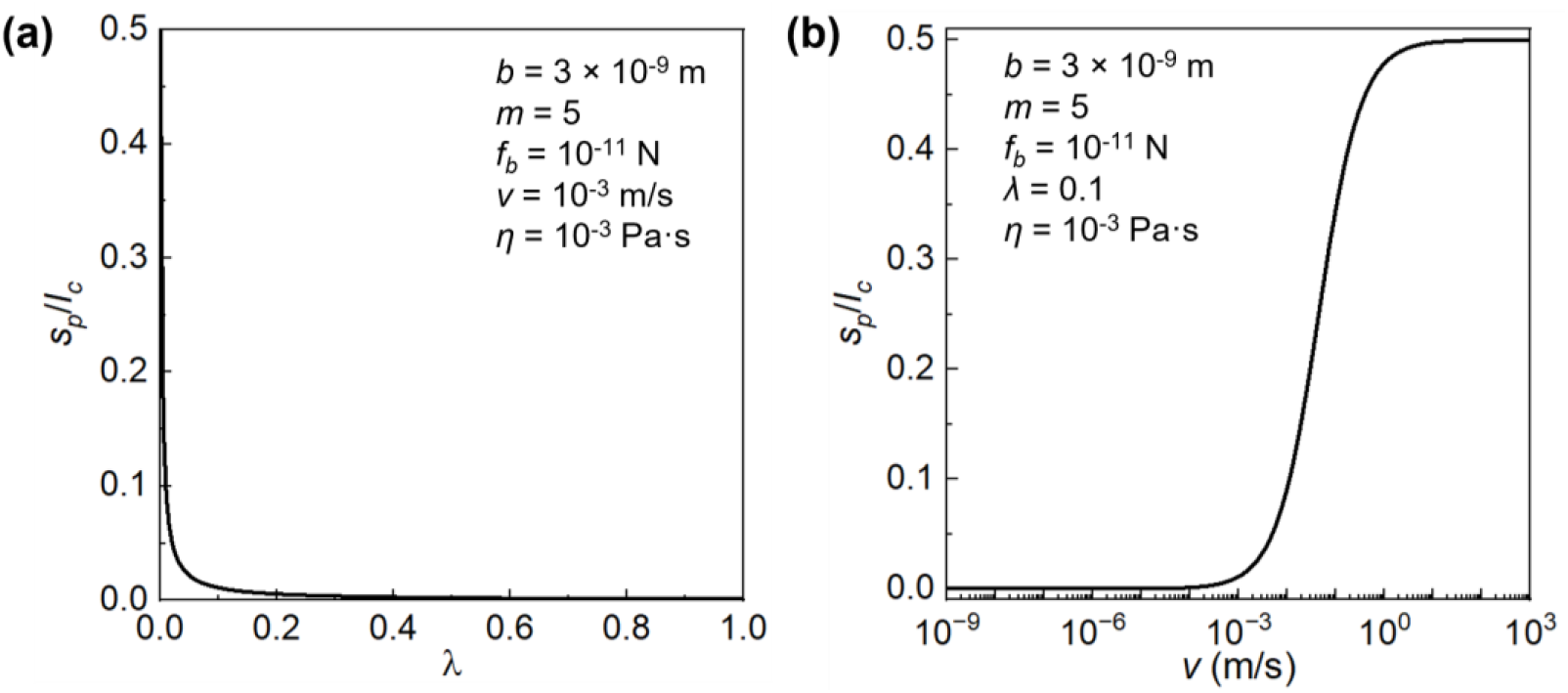
Chain penetration depth for equal force pullout. Penetration depth versus **(a)** bond formation parameter, and **(b)** pullout velocity.

Rewriting Eq. (19) to include all chains, the stress for chain pullout from the anionic network is

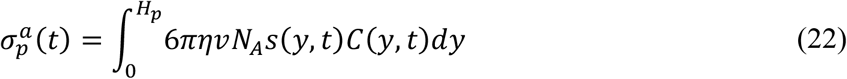

and from the e-GLUE network is

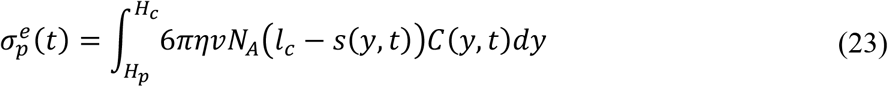

### 3.6. Debonding stress

During debonding, the polycations are pulled out from both networks, depending on how deeply they penetrate the anionic network. For chain heads that reside within *H*_*p*_, they are subject to bond breaking and pullout from the anionic network. For chain heads that reside between *H*_*p*_ and *H*_*c*_, they are pulled out from the e-GLUE network. By combining Eq.(18), (22), and (23), the adhesion strength is

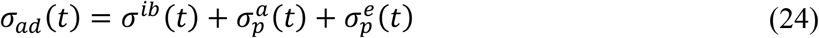

## 4. Model results

### 4.1. Parameters used in the calculation

For a polycation, e.g., PDAC, the monomer length *l*_*m*_ = 0.55 nm and the Kuhn length *b* = 3 nm (ref^41^). Each Kuhn segment contains approximately 5 monomers and therefore carries 5 cations, *m* = 5. The molecular weight (number-average Mw) of the chain is fixed at 100 kDa (*n* = 124, given the monomer Mw is 161.5 Da) and is varied from 10 kDa to 1,000 kDa to study the Mw effect (*n* = 12 to 1,238). The cation concentration is fixed at 0.691 M when studying the effect of other parameters, and is varied from 0.1 M to 5 M when studying its own effect on ionic bond formation and adhesion. The zero-field friction coefficient is fixed at 10^−5^ kg/s when studying the effect of other parameters, and is varied from 5 × 10^−6^ to 10^−4^ kg/s when studying its own effect on chain migration, ionic bond formation, and adhesion. These values are roughly 10^−15^ m^2^/s to 10^−20^ m^2^/s for chain diffusivity, which is in a reasonable range from reported values^18^. The chain activation length is set as the Kuhn length. For the anionic network, the anion concentration is fixed at 0.085 M. The electric field is fixed at 5,000 V/m when studying the effect of other parameters, and is varied between 0 V/m and 7,500 V/m when studying its own effect on chain migration, ionic bond formation, and adhesion. The applied duration of the electric field is less than 15 s. The ionic bond formation parameter is set as 0.1 (ref^42^), and the force to break an ionic bond is 10 pN (ref^17^). The viscosity of water is 0.001 Pa·s. The pulling speed is fixed at 0.001 m/s.

### 4.2. Polycation distribution in electrophoresis

Of particular interest is the chain migration kinetics under an electric field. The chains migrate deeper into the anionic network over time, either by pure diffusion in the absence of an electric field or by electrophoresis. The applied electric field can enhance migration, but the effect is not strong (Fig. 6a). For example, at 15 s, under no electric field, the chains penetrate about 0.32*l*_*e*_, in contrast to penetrating 0.37*l*_*e*_ with *E* = 7,500 V/m. Such a partial chain penetration in the anionic network is beneficial to bridge the interface for adhesion. We next observe chain distribution at 6 s. The chain concentration decreases with deeper migration. Though the electric field can increase it, the increment is small (Fig. 6b). We further obtain the cation concentration distribution by integrating chain concentration curves at each location using Eq. (9) and show that the concentration increases with the electric field (Fig. 6c). We also note that the concentrations at 6 s is about 0.01 M, one order of magnitude lower than the anion concentration (0.085 M), suggesting that the number of cations determines the number of ionic bonds, and these ionic bonds have not reached saturation.

**Fig. 6.**
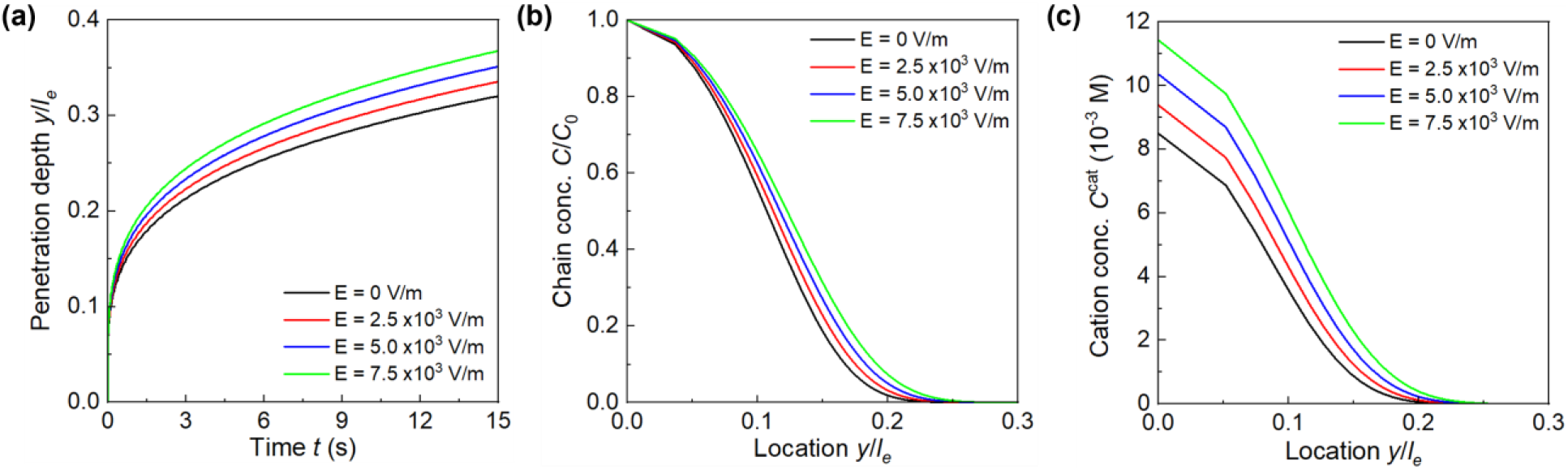
Effect of electric field on chain migration. **(a)** Chain penetration depth versus time. Distributions of **(b)** chain and **(c)** cation concentrations in the anionic network under various electric fields at 6 s.

We next study the chain length effect on migration. A longer chain is more difficult to mix with a hydrogel as the mixing is less favorable than a small molecule, and the osmotic pressure generated by the hydrogel network favors demixing^43^. A longer chain also experiences more resistance during migration, e.g., ionic bonding and chain entanglement, leading to slower migration compared to a short chain (Fig. 7a). A short chain with Mw = 10 kDa can migrate beyond its size about 1.15 s, and a medium chain with Mw = 50 and 100 kDa can still migrate 0.57*l*_*e*_ and 0.35*l*_*e*_ at 15 s. In contrast, a long chain with Mw = 1,000 kDa just penetrates about 0.14*l*_*e*_. In addition, a long chain exhibits a significantly lower concentration compared to a shorter chain at the same location. Though the difference is small near the interface, it becomes prominent farther away from the interface (Fig. 7b). In particular, the long chains with Mw = 1,000 kDa are largely localized near the interface (*y* < 0.1*l*_*e*_), whereas the short chains with Mw = 10 kDa have a broad distribution across *l*_*e*_. The calculated cation concentration further shows that the cation concentration with Mw = 10 kDa already exceeds the anion concentration with *y* < 0.7*l*_*e*_, yet the chains with a larger Mw are still below it (Fig. 7c).

**Fig. 7.**
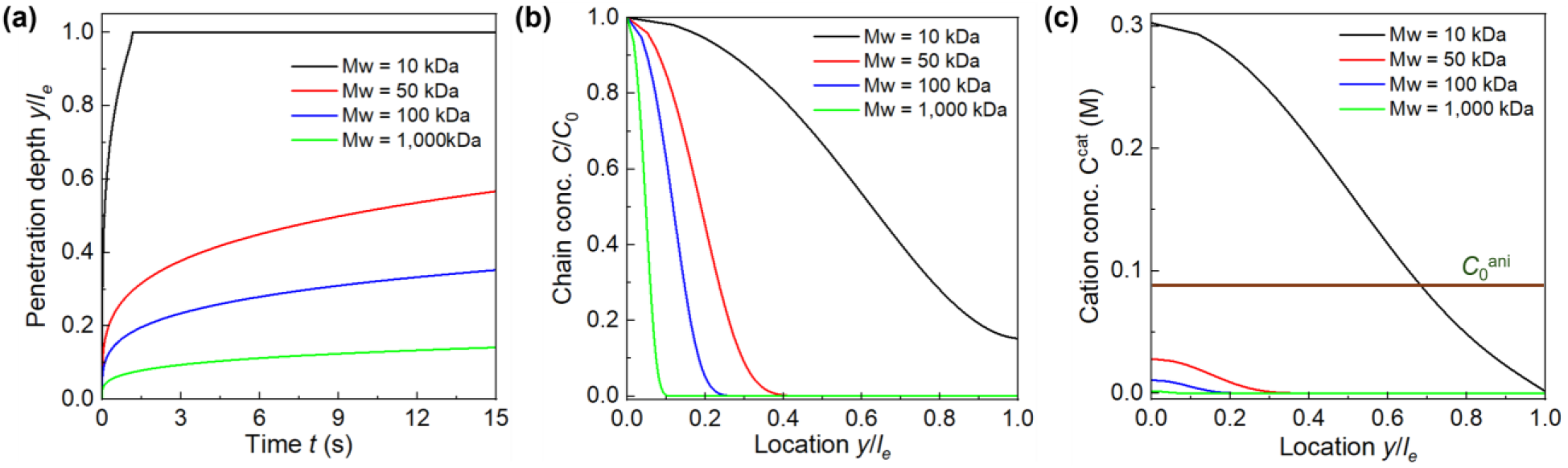
Effect of molecular weight on chain migration. **(a)** Chain penetration depth versus time. Distributions of **(b)** chain and **(c)** cation concentration in the anionic network under various molecular weights at 6 s.

Types of cations and anions determine the strength of their ionic pairs^44, 45^, which reflects the migration stickiness of polycations. The zero-field friction coefficient ξ_0_ describes such stickiness. It is clear that a large ξ_0_ retards migration, and the electric field can assist in migration, though its effect is less strong than the effect of molecular weight (Fig. 8a). For example, when ξ_0_ = 10^−4^ kg/s, the chains only migrate a tiny distance of about 0.145*l*_*e*_ without the electric field, and can just migrate to 0.164*l*_*e*_ with *E* = 7,500 V/m. Similarly, a bigger ξ_0_ reduces both the chain (Fig. 8b) and cation concentration (Fig. 8c) more significantly.

**Fig. 8.**
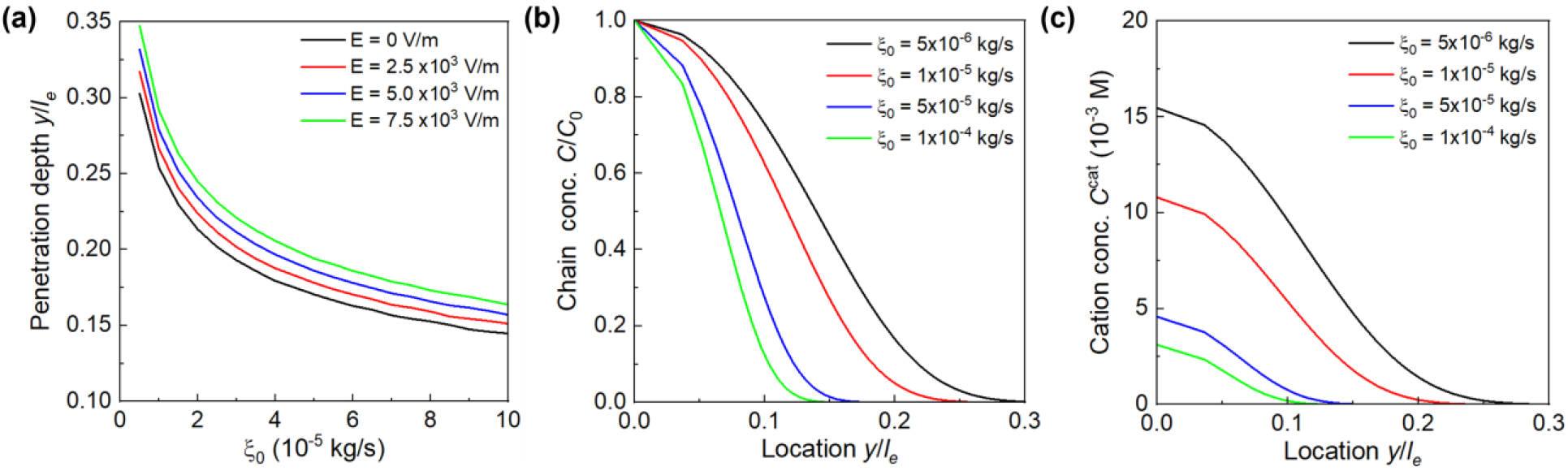
Effect of friction coefficient on chain migration. **(a)** Chain penetration depth versus friction coefficient under various electric fields. Distributions of **(b)** chain and **(c)** cation concentration in the anionic network under various friction coefficients at 6 s.

### 4.3. Formation of ionic bonds in electrophoresis

Having obtained the spatial distribution of cation concentration with time, we next determine the concentration of the total formed ionic bonds with time. Here, we consider ideal bond formation without adjusting it with the bond formation parameter (*λ* = 1). Our calculation confirms that ionic bonds accumulate over time and increase with the electric field, but do not reach saturation in 15 s (Fig. 9a), because the cation concentration is lower than the anion concentration (Fig. 6c). We vary molecular weights and observe that the ionic bond concentration reduces significantly with chains with Mw > 50 kDa (Fig. 9b). For any chain length, a bigger friction reduces ionic bond concentration, and the reduction becomes bigger with longer time (Fig. 9c). Increasing *C*_0_^*cat*^ allows more chains to enter the anionic network at the same time to generate more ionic bonds (Fig. 9d). For example, increasing *C*_0_^*cat*^ from 0.5 M to 5 M can form almost 10 times more ionic bonds.

**Fig. 9.**
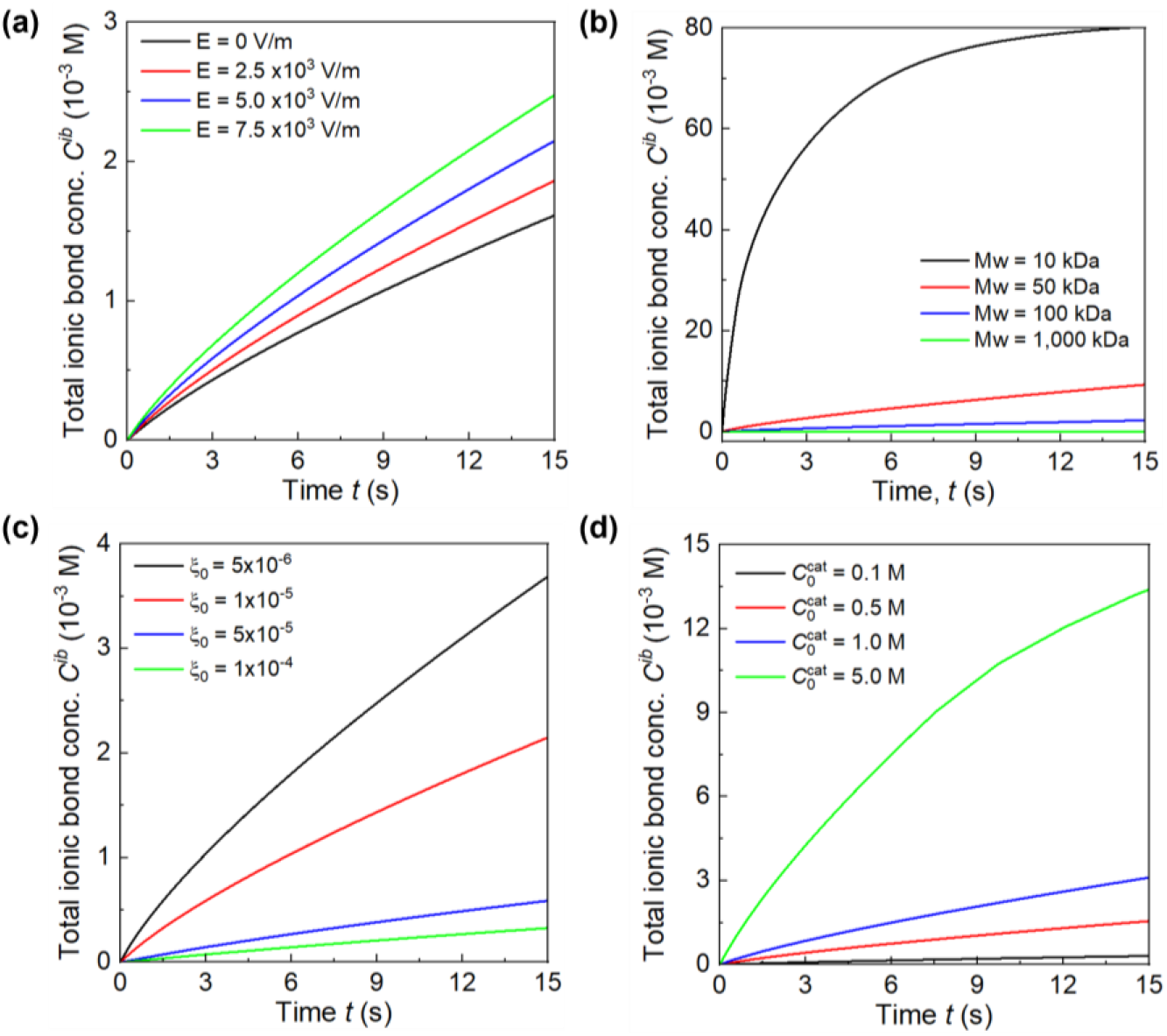
Formation of total ionic bonds with time. Total ionic bond concentration versus time with **(a)** electric field, **(b)** molecular weight, **(c)** friction coefficient, and **(d)** cation concentration.

We also compare the concentrations of the total formation of ionic bonds and the ionic bonds to be breaking (Fig. 10). Given typical chain parameters, the equal chain pullout depth *s*_*p*_ = 0.011*l*_*c*_, or *H*_*p*_ = 0.105*l*_*e*_ (Fig. 5a). In the early migration within 0.7 s, the chains just penetrate the anionic network with a depth of about 0.1*l*_*e*_ (Fig. 6a), consequently, the total formation of ionic bonds is the same as those to be broken during debonding. As chains penetrate deeper beyond *H*_*p*_, these ionic bonds within *H*_*p*_ that can originally break during debonding, nonetheless, no longer break, leading to a gradually enlarged difference between the two concentrations.

**Fig. 10.**
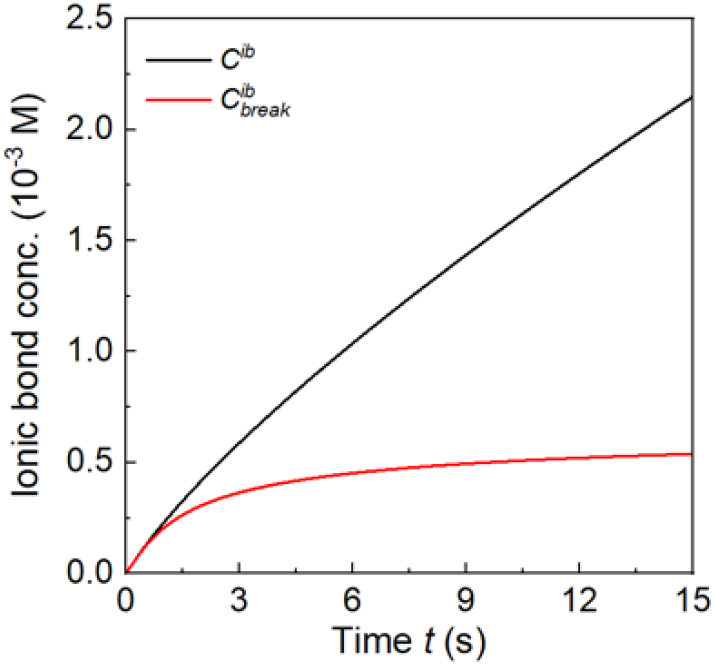
Comparison of the total formation of ionic bonds and ionic bonds to be broken.

### 4.4. Adhesion strength in electrophoresis

We first calculate the individual contribution of ionic bond breaking, chain pullout from the e-GLUE, and chain pullout from the anionic network to the adhesion strength. The result shows that the adhesion strength increases with time, in which the stress for breaking ionic bonds contributes dominantly and that for chain pullout from the e-GLUE contributes modestly, while the stress for chain pullout from the anionic network is negligible (Fig. 11a). In particular, the breaking stress of ionic bonds and the adhesion strength are almost identical in the first 1 s, and then the chain pullout stress from the e-GLUE starts to play a role. This behavior is consistent with Fig. 10, where the shallow penetration breaks the ionic bonds and pulls out the chains from the anionic network during debonding. Because the chain pullout distance is tiny, ∼0.105*l*_*e*_, the pullout force is small, and the ionic bond breaking stress determines the adhesion strength. As more chains pass through *H*_*p*_, the debonding starts to pull out more chains from the e-GLUE, leading to its increased contribution. Adhesion strength increases marginally with the electric field (Fig. 11b).

**Fig. 11.**
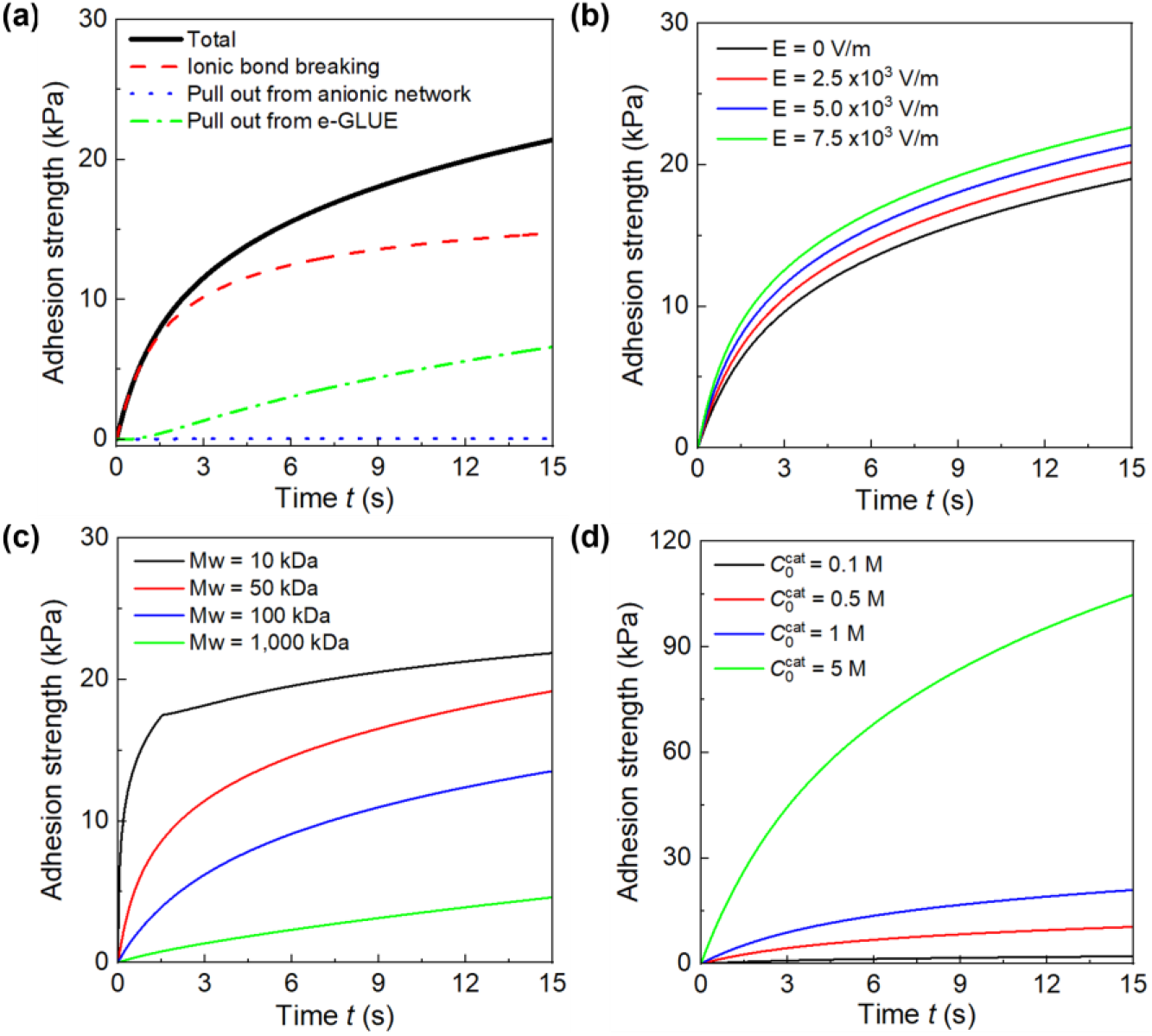
Formation of adhesion with time. **(a)** The contributions of ionic bond breaking and chain pullout to the adhesion strength. The adhesion strength versus time under various **(b)** electric fields, **(c)** molecular weights, and **(d)** cation concentrations.

At 15 s, the adhesion strength is about 19 kPa with *E* = 0, and rises to 22.6 kPa with *E* = 7,500 V/m. We also observe that short chains with Mw = 10 kDa boost adhesion rapidly in 1.5 s and increase much more slowly afterward, whereas other chains continuously develop adhesion strength with time (Fig. 11c). The longer chains are likely to surpass the adhesion strength of short chains, provided a longer time. This result suggests that a short chain can quickly enable adhesion, but penetration is superficial; a long chain can penetrate deeply, but it takes time to build adhesion. Finally, increasing cation concentration strengthens adhesion proportionally (Fig. 11d). For example, concentrating cations by 10 times increases the adhesion strength by 10 times. Here, we point out that the same cation concentration can be achieved by either using short chains at a high concentration or long chains at a low concentration, which yields distinct outcomes: the former can enable rapid but shallow adhesion, whereas the latter can generate slow but deep adhesion.

## 5. Experiments

We conduct electroadhesion experiments to validate the theoretical model. We synthesize a polyacrylamide (PAAm) hydrogel with mobile PDAC as the e-GLUE. We slightly add chitosan and sodium tripolyphosphate (TPP) to enhance stiffness and handleability. We synthesize a PAAm hydrogel with an anionic alginate network as the mucosa. We measure adhesion strength with time, electric field, and chain length.

### 5.1. Materials

Acrylamide (AAm; A8887), alginate (A2033), PDAC chains of Mw = 400 kDa – 500 kDa (409030), Mw = 200 kDa – 350 kDa (409022), and Mw <100 kDa (522376), chitosan (448877), covalent crosslinker *N, N*′-methylenebis(acrylamide) (MBAA; M7279), ionic crosslinker calcium sulfate (CaSO4; 255548) and sodium tripolyphosphate (TPP; 238503), initiator ammonium persulfate (APS; A3678) and initiator accelerator tetramethylethylenediamine (TEMED; T7024) were purchased from Sigma-Aldrich.

### 5.2. Synthesis of the e-GLUE and the anionic hydrogel

The e-GLUE was synthesized as follows. Chitosan and AAm were first dissolved in acetic acid solution (pH = 5) at 2 and 12 wt%, respectively, and stirred to obtain a clear solution. After degassing, 5 ml of the precursor solution, 36 μl of 2 wt % MBAA, and 8 μl TEMED were mixed in one syringe, and 5 ml of 20 wt% PDAC, 226 μl of 0.27 M APS, and 52.5 μl of 0.75 M TPP were mixed in another syringe. The two syringes were mixed, and the mixture was cast in a glass mold at room temperature overnight to complete polymerization.

The anionic hydrogel was synthesized as follows. Sodium alginate and AAm were dissolved in DI water at 2 and 12 wt% and stirred overnight. After degassing, a 10 ml solution was mixed with 36 μl of 2 wt% MBAA and 8 μl TEMED in one syringe. Separately, 226 μl of 0.27 M APS and 191 μl of 0.75 M CaSO_4_ slurry were mixed in another syringe. The two syringes were mixed, cast in a glass mold, and polymerized overnight at room temperature.

### 5.3. Adhesion test

The e-GLUE was placed on top of the anionic hydrogel with a contact area of 4 cm^2^ (Fig. 12a), which was then sandwiched between two parallel electrodes made of stainless steel to complete the electroadhesion setup (Fig. 12b). A voltage was supplied across the electrodes in 15 s to minimize electrochemical reactions, though mild electrode redox reactions and water electrolysis still occurred. Adhesion strength was measured by the probe-pull test after turning off the voltage (Fig. 12c). The top e-GLUE and the bottom anionic hydrogel were bonded to rigid PET films with Krazy glue to restrict deformation. A tensile force was applied to pull the two hydrogels apart at a rate of 1 mm/s using a tensile machine (Mark-10 ES20) equipped with a digital force gauge (Mark-10 M4-05). The tensile force as a function of time was recorded, and the adhesive strength was calculated by normalizing the peak tensile force by the contact area.

**Fig. 12.**
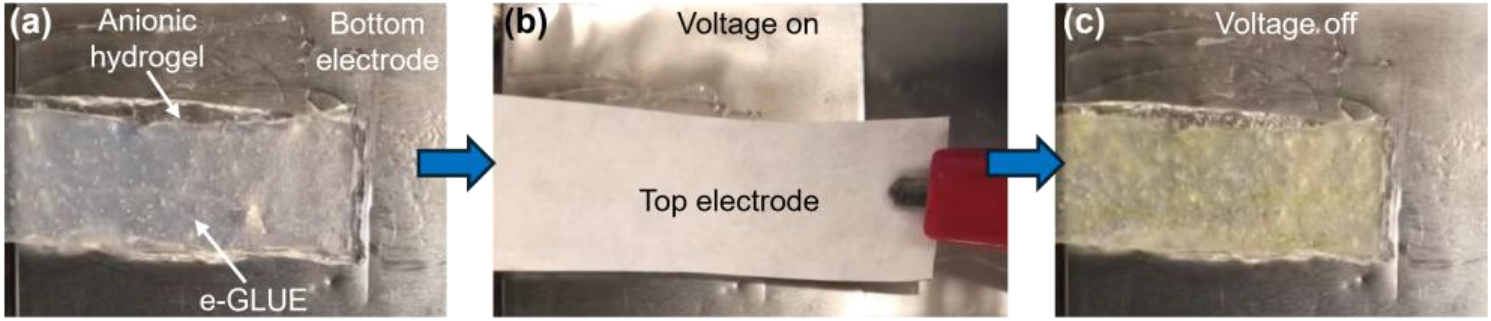
Electroadhesion experiments. **(a)** Test setup. **(b)** Apply voltage. **(c)** Adhesion forms.

### 5.4. Experimental results

We first seek to determine ξ_0_ and *L* in the model by fitting the experimental data. The PDAC chains in the e-GLUE have a wide distribution of Mw of 200 – 350 kDa. We therefore choose the average Mw = 275 kDa in the model. We applied *E* = 4,000 V/m and measured the adhesion strength with time. The data show that the adhesion strength increases with time, from 2.5 kPa up to 17 kPa in 15 s (Fig. 13a). A large variation is observed at a longer time, possibly due to voltage-induced hydrogel deterioration and water electrolysis that degrade adhesion. By fitting the data, we find ξ_0_ = 3 × 10^−5^ kg/s and *L* = 8.9 nm. The ξ_0_ value falls into a common range from reported values^18^, and the *L* value is on the order of a Kuhn length, both of which are consistent with our model setup. We then use these fitting parameters to predict adhesion strength as a function of electric field, which exhibits nice agreement (Fig. 13b). The adhesion strength enhances from 1.3 kPa without the electric field to 24 kPa with *E* = 6,667 V/m. We lastly vary PDAC chains with a higher Mw of 400 kDa – 500 kDa and a lower Mw < 100 kDa, and observe that the adhesion strength is higher with shorter chains. To compare with our model prediction, we implement average Mw = 450 kDa and Mw = 100 kDa in the model. Because Kuhn segment friction does not depend on chain length, we fix ξ_0_ and fit *L* by the experimental data, which gives *L* = 9.1 nm for Mw < 100 kDa and 7.4 nm for Mw of 400 kDa – 500 kDa (Fig. 13c). These activation lengths are similar, on the order of a Kuhn length, and independent of chain length. This result is expected, as overcoming friction over a Kuhn distance is local rather than global.

**Fig. 13.**
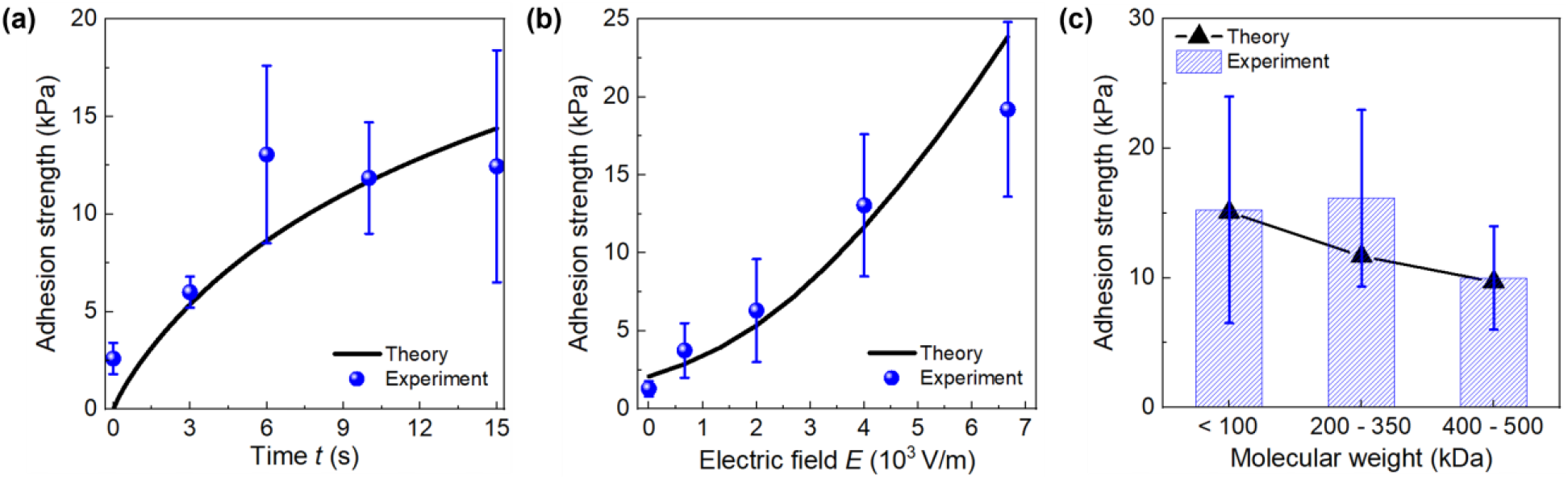
Electroadhesion tests. Compare adhesion strength versus **(a)** time, **(b)** electric field, and **(c)** molecular weight between experimental data and model predictions.

## 6. Discussion

The model developed in this work enables rational, quantitative design and control of the electrical field, applied duration, chain length, and cation concentration for e-GLUE mucosal adhesion. The goal is to establish mucosal adhesion with high adhesion strength, a short application time, deep chain penetration, and minimal impact on mucosal physiology. In particular, adhesion strength needs to match the mechanical demands of the target mucosal environment. For example, the GI tract is subject to continuous peristalsis, luminal flow, and cyclic deformation, generating stresses in the kPa range under strong contraction^46, 47^. To ensure stable attachment, the adhesion strength should exceed these forces with an appropriate safety margin, e.g., tens of kPa, whereas achieving hundreds of kPa may not be necessary.

The electric field is a critical parameter to enhance chain migration and mixing with the mucosa to establish strong adhesion, yet its magnitude must be carefully tailored to ensure tissue safety. A high electric field (e.g., > 20,000 V/m) generates a strong current to induce undesirable heat and electrochemical reactions^48, 49^, which are detrimental for sensitive mucosal tissues such as the colon ^7^. Conversely, a low electric field (e.g., < 4,000 V/m) results in weak electrophoresis, which may take a longer time (> 6 s) to reach an adhesion strength of tens of kPa. In our experiments, an electric field of 4,000-7,000 V/m generating an adhesion strength of 10 kPa in 6 s is both tissue-safe and adhesion-effective (Fig. 13b). Another possible strategy is to use pulsed signals of a high electric field > 20,000 V/m to allow intermittent heat dissipation while maintaining chain electrophoresis to build up adhesion.

A longer electric field application generally leads to stronger adhesion, and the increment depends on chain length. Short chains rapidly increase adhesion to a relatively high level in the first few seconds, yet the further increase is much slower. Such a few-second application may be ideal in practice. By contrast, long chains require more time to develop adhesion, though experiments show that the adhesion strength is almost similar after 6 s (Fig. 13a). Consequently, an application duration longer than 6 s may not be necessary, and the optimal duration depends on the chain length and the target adhesion level.

Chain length not only influences migration kinetics but also the critical penetration depth. Short chains form adhesion quickly but penetrate only superficially, inhibiting deep adhesion. By contrast, long chains form adhesion slowly but can migrate deeper to improve adhesion retention. Therefore, long chains, driven by a pulsed high electric field, can enable deep penetration. It is ideal to migrate chains across the mucus and epithelium layers of 100 nm thick to anchor them to more stable tissue compartments, but finding chains with the required large molecular weight may be challenging.

Increasing cation concentration is straightforward to enhance adhesion (Fig. 11d). During the synthesis of e-GLUE, it is key to dissolve polycations near their solubility limit. Because chains with a high molecular weight typically have lower solubility than those with a low molecular weight, the molecular weight selection and its associated solubility need to be investigated and balanced to achieve the maximum cation concentration while maintaining processability.

Taken together, these considerations suggest that an ideal e-GLUE mucosal adhesion may employ a hydrogel containing long-chain polycations near their solubility limit and operate under a consistent, pulsed, high-electric-field to generate strong, deep adhesion. Another strategy is to use a mixture of short- and long-chains in a specific ratio, in which short chains establish rapid adhesion in the initial few seconds, while long chains gradually migrate deeper into the mucosa and form deeper adhesion over time, enabling both rapid and long-retention adhesion.

## 7. Conclusion

We formulate a theoretical model to describe the sticky electrophoresis, ionic complexation, and chain entanglement of polycations for the polycation interfacial bridging electroadhesion. We quantitatively derive the migration kinetics, ionic bond formation, and adhesion strength, with polycation chain length, friction coefficient, and cation concentration, as well as electric field and duration. Calculations show that the stress for ionic bond breaking is a major contributor to the adhesion strength compared to that for chain pullout. Adhesion strength increases with stronger electric fields, longer application duration, shorter chains, and higher cation concentration. We further perform electroadhesion tests, and the measured adhesion strength is in good agreement with the values predicted by our model. Using this model, we lastly suggest potential design approaches for the e-GLUE.

## Acknowledgments

J.Y. acknowledges support from the National Science Foundation CAREER Award (DMR-2440130) and the James Nichols Heald Startup Award from Worcester Polytechnic Institute. B.Y. acknowledges support from the University of Texas Southwestern Medical Center Startup Fund, American Cancer Society Institutional Research Grant (IRG-24-1322075-19-IRG), and UTSW Cancer Center Support Grant (P30CA142543). K.Y. acknowledges support from the College of Engineering and Computer Science at Syracuse University. S.Y. acknowledges support from the National Science Foundation Future Manufacturing Research Grant (FMRG, #CMMI-2037097). The authors acknowledge the helpful discussion with Dr. An Xin.

